# Dissection of Poly(A)-binding protein (PABPC) cellular function using degron-mediated depletion with replacement

**DOI:** 10.64898/2026.06.05.728833

**Authors:** Ryan Y Muller, Kiara X Wang, David P Bartel

## Abstract

Cytoplasmic poly(A)-binding proteins (PABPCs) are essential and highly abundant regulators of mRNA stability and translation, but their cellular functions have been difficult to dissect due to slow turnover and the lethality observed upon loss of both major paralogs, PABPC1 and PABPC4. To enable structure–function analysis of PABPC in human cells, we developed a protein-replacement platform that couples rapid auxin-inducible degradation of endogenous PABPC1 and PABPC4 with doxycycline-controlled expression of engineered PABPC variants. This approach enables acute removal of native PABPC and real-time assessment of how specific domains, paralogs, and sequence alterations support cellular fitness, shape transcriptome profiles, and regulate poly(A)-tail length. Using this system, we show that the RRM4 domain of PABPC1 is essential for growth, whereas post-translationally modified lysines within RRM4 are individually dispensable. Paralogs and variants with heterologous RRM4 domains vary in their ability to substitute for PABPC1, revealing functional divergence among PABPCs. Transcriptome profiling identifies variant-specific regulatory signatures, and dose-controlled rescue further delineates the relationship between PABPC variant abundance and global poly(A)-tail lengths in vivo. Together, this platform provides a generalizable strategy for dissecting PABPC biology in the cellular context using rationally designed variants.

## Introduction

Cytoplasmic poly(A)-binding proteins (PABPCs) are essential regulators of mRNA stability, translation initiation, and translation termination^1,2^. In mammals, the ubiquitously expressed PABPC1 and its paralog PABPC4 bind polyadenylated transcripts via their four RNA recognition motifs (RRMs) and recruit regulatory factors through their C-terminal Methionine-Leucine-Leucine-Glutamate (MLLE) domain^3,4^. Additionally, PABPC can further regulate transcript expression with interactions involving other regions of the protein, such as the interactions with eukaryotic initiation factor 4G (eIF4G)^5,6^. The sum of PABPC interactions result in a complex and nuanced landscape of PABPC-mediated gene regulation. Biochemical approaches have contributed much to our understanding of interactions within this regulatory landscape, but are not able to determine how these interactions contribute to regulation in the cellular context.

Studies to characterize the cellular function of PABPC would benefit from an ability to replace endogenous PABPCs with designed variants. PABPC1 and PABPC4 present a unique experimental challenge in this regard, owing to their high abundance and essentiality for viability. Traditional RNAi- or CRISPR-based depletion strategies must contend with slow turnover rates, incomplete knockdown, and double-knockout inviability. Thus, we and others have used the auxin-inducible degron (AID) system^7,8^ for rapid and conditional degradation of endogenous PABPC1 and PABPC4. The success of this approach for enabling acute loss-of-function experiments implies that it might be adapted for systematic protein replacement, in which endogenous PABPCs are rapidly removed and functionally substituted with engineered variants.

Here, we establish a degron-mediated protein replacement platform for dissecting PABPC function in human cells. In this approach, endogenous PABPC1 and PABPC4 are tagged with an mAID degron and degraded upon addition of indole-3-acetic acid (IAA, an auxin analog), while a doxycycline-inducible PABPC rescue variant is expressed from a stably integrated construct. By combining rapid depletion with tunable rescue expression, this method allows direct assessment of how specific PABPC domains, paralogs, and engineered sequence variants support cell growth, shape the transcriptome, and influence mRNA poly(A)-tail length.

## Results

### Endogenous PABPC can be replaced with PABPC variants

To study the role of PABPC1 and PABPC4 in translational regulation, our lab previously used an inducible AID endogenous degron tag for rapid degradation of PABPC1 and PABPC4^9^. We reasoned that this same depletion strategy might help further interrogate PABPC1 function in a cellular context, through rapid depletion of the endogenous protein and rescue with a designed PABPC variant. To implement this strategy, polyclonal cell lines were generated, in which doxycycline (dox)-inducible PABPC rescue variants were integrated into a parent cell line that expressed dox-inducible OsTir1 and had mAID tags on all endogenous PABPC1 and PABPC4 alleles (fig 1A). Treatment with 1µg/mL dox, followed by 0.5mM (IAA), was sufficient to degrade endogenous PABPC1 and PABPC4 while also inducing expression from the integrated construct (fig 1B). Testing this approach with several PABPC variants (fig 1C) yielded similar levels of rescue expression across PABPC variants (fig 1D).

**Figure 1.**
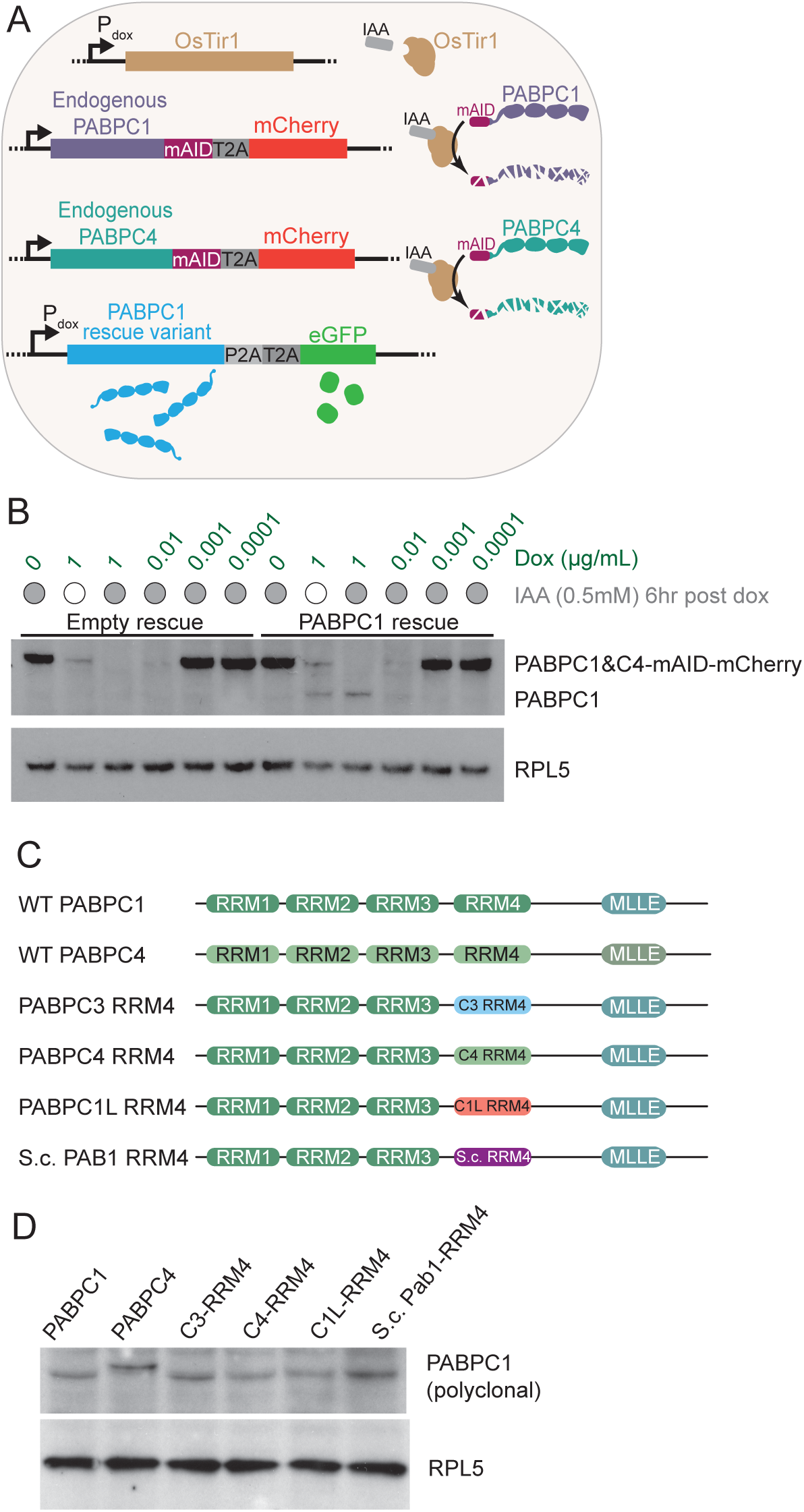
Endogenous PABPC can be replaced with PABPC variants. (A) Schematic of the degron-mediated protein-replacement strategy. Parental HCT116 cells express the OsTIR1 F-box protein and harbor mAID–mCherry–T2A tags on both endogenous PABPC1 and endogenous PABPC4. Addition of doxycycline (Dox) induces expression of OsTir1 and a PABPC rescue variant from a stably-integrated constructs, while treatment with indole-3-acetic acid (IAA) triggers rapid degradation of endogenous mAID-tagged PABPC1 and PABPC4. (B) Immunoblot validation of endogenous PABPC1 and PABPC4 degradation and rescue variant expression following treatment with indicated concentrations of doxycycline and 0.5 mM IAA. IAA was provided 6 h after doxycycline, and cell lysates were collected 24 h after doxycycline addition. RPL5 served as a loading control. (C) Domain organization of wild-type PABPC1 and RRM4 replacement variants, including human paralog PABPC4 and variants that replaced the PABPC1 RRM4 with the RRM4 domain of either PABPC3, PABPC4, PABPC1L, or yeast PAB1 RRM4. (D) Immunoblots showing that PABPC rescue variants have similar expression levels after PABPC replacement. Lysates were collected 21 days after doxycycline treatment.

### Growth phenotypes indicate function of protein domains within PABPC1

After using this approach to replace endogenous PABPC with PABPC variants, we performed growth assays that sought to provide insights regarding how domains within PABPC1 functioned in a cellular context. In these assays, cells were counted and split every 6–7 days to monitor growth rate.

Yeast PABPC1 is reported to straddle the boundary between the 3’ UTR and the polyA tail, placing RRM4 of PABPC1 on the 3’ UTR and implying that RRM4 might be more promiscuous in its RNA-binding specificity^10^. We used our PABPC replacement approach to perturb RRM4 and observe the effects on growth. As expected, PABPC1 and PABPC4 expression were both sufficient to rescue growth upon endogenous PABPC depletion, whereas an empty construct failed to rescue growth (fig 2A). Deletion of RRM4 also was not viable, nor was a Y297A mutation within RRM4 that was designed to interrupt RNA binding of RRM4, modeled after the F367A mutation in *S. pombe* Pab1 reported to ablate RRM4 binding^10^.

**Figure 2.**
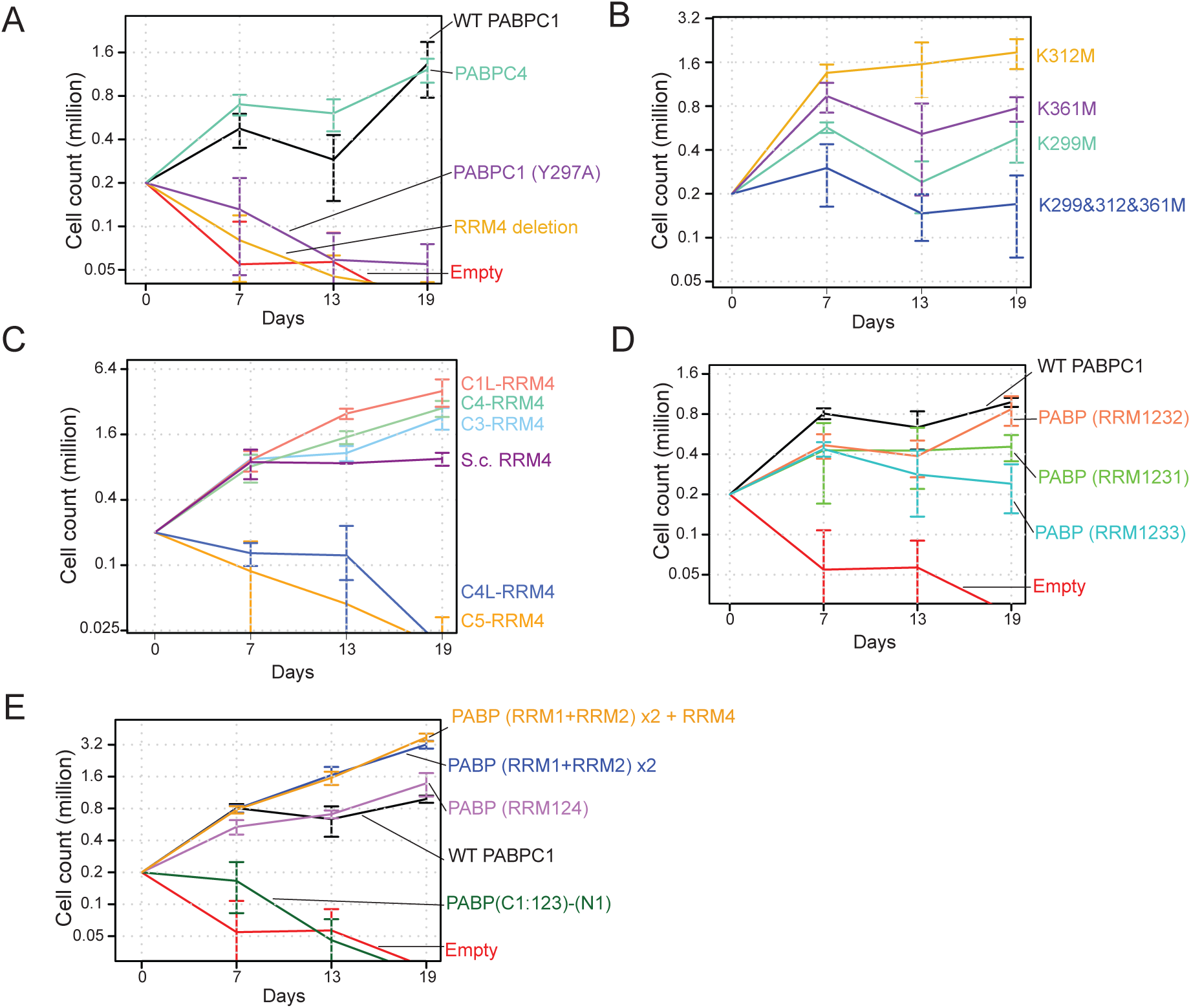
Growth phenotypes dissect function of protein domains within PABPC1. (A) Growth curves following endogenous PABPC1 and PABPC4 depletion and rescue with either wild-type or mutant PABPC variants. PABPC1 and PABPC4 fully rescued growth, whereas an empty vector, deletion of RRM4, or a Y297A point mutation in RRM4 failed to support growth. Cells were seeded at 0.2 × 10⁶ and split 1:3 on days 7 and 13. (B) Growth assays testing lysine-to-methionine substitutions within RRM4 (K299M, K312M, K361M, and triple K299/312/361M) (C) Growth assays testing rescue of variants in which RRM4 of PABPC1 is replaced with RRM4 domains from other human PABPC paralogs (PABPC3, PABPC4, PABPC1L, PABPC4L, or PABPC5) or from yeast PAB1. (D) Growth assays replacing RRM4 with other PABPC1 RRM domains (RRM1, RRM2, RRM3) or (E) Growth assays testing altered numbers of domains and a PABP1212 variant.

With the observation that RRM4 was necessary to rescue growth, we interrogated perturbations within RRM4. Of the previously identified post-translational modifications in PABPC1, four occur on lysines within RRM4^11^. These modifications include K299 methylation, K312 acetylation or dimethylation, and K361 dimethylation. Given that lysine residues are often important for RNA binding, we wondered if lysine modifications might play a role in modulating RNA-binding activity. We mutated each lysine to methionine, which is similar in size to lysine but is not positively charged and cannot be post-translationally modified. Mutations in any of the three lysines supported growth (fig 2B). Mass spectrometry of purified SBP-tagged PABPC1 ectopically expressed in HEK cells detected modification K299, but not of the other three modifications previously reported within RRM4 (fig S1), suggesting that the other modifications might occur more rarely. The K299 mutation also had the largest impact of the three single K-to-M substitutions in our growth assay (fig 2B). The triple substitution had a stronger impact than any of the single K-to-M substitutions, but still supported cell growth (fig 2B).

The development of this approach also provided the opportunity to investigate how the RRM4 domains of different PABPC proteins might differ from each other. The RRM4 domain of PABPC1 was substituted with the RRM4 of each PABPC1 ortholog expressed in humans and yeast, and then these fusion proteins were tested for their ability to complement loss of endogenous PABPC1/4 (fig 2C). Most RRM4 domains were able to rescue cell growth. The exceptions were the RRM4 domains from PABPC5 and PABPC4L. Based on GTEx bulk tissue expression measurements, PABPC5 and PABPC4L are the lowest expressed among human PABPC paralogs across tissues. Across tissues, the expression of *PABPC5* is highest in ovaries, while *PABPC4L* is highest in uterus. These two human paralogs also diverge the most from human PABPC1 at both the nucleotide and protein sequence level.

Previous work shows that a truncated PABPC1 containing RRM3 and RRM4 has relaxed preferences for polyA sequence compared to RRM1 and RRM2^12^. From our growth rescue observations, we wondered if RRM4 functions within PABPC1 through potential sequence preferences within the 3’ UTR or if general RNA affinity afforded by any PABPC domain would be sufficient to rescue growth. Accordingly, the RRM4 of PABPC1 was replaced with each other RRM domain in PABPC1 (fig 2D). Each substitution rescued growth, implying that the generic activity of any of these RRM domains was sufficient.

With the understanding that specific sequence preferences of RRM4 were not required for growth rescue, we tested an additional PABPC variant comprised of repeat of domains RRM1 and RRM2, referred to as PABP1212 (fig 2E). This PABPC variant was predicted to have a strong preference for pure polyA sequences, because RRM1 and RRM2 have substantial stringency for polyA sequence compared to RRM3 and RRM4^13,14^. PABP1212 rescued growth, indicating that any sequence preferences conferred by RRM3 and RRM4 are not necessary for cell growth. Additional PABPC variants containing either three (RRM124), four (RRM1212), or five (RRM12124) domains each rescued growth (fig 2E), indicating the PABPC1 footprint size on RNA is also not a necessary component of PABPC1 function in our cultured cells.

### PABPC variants influence the cellular transcriptome

To assess the transcriptome-wide consequences of PABPC1 domain substitutions, we performed RNA-seq under conditions in which endogenous PABPC1/4 was depleted and replaced with individual PABPC constructs (fig 3A). Our growth assays showed that some PABPC variants rescued growth, whereas others did not. The transcriptomes of rescuing PABPC variants were sequenced 25 days after drug treatment, whereas the transcriptomes of non-rescuing PABPC variants were sequenced 48 h after doxycycline and IAA treatment, while cells were still alive (fig 3A). Cells with the control PABPC1 rescue were sequenced at both 48h and 25 days. PCA analysis was used to compare transcriptomes across replacements with different PABPC variants. When comparing all the PABPC variants that were sequenced after 48 h, those lacking RRM4 or containing heterologous RRM4 domains (human PABPC5, PABPC4L, or yeast PABPCs) segregated into PCA clusters away from PABPC1 48 h rescue, indicating large-scale transcriptional divergence from the wild-type rescue profile (fig S2A). Removing RRM4 of PABPC1 was more similar to a lack of PABPC1 (empty) than it was to replacement of RRM4 with that of PABPC4L or PABPC5 (fig S2A). These results suggested the RRM4 domains of PABPC4L and PABPC5 retain some function as RNA-binding domains, although with activity that fails to rescue growth.

**Figure 3.**
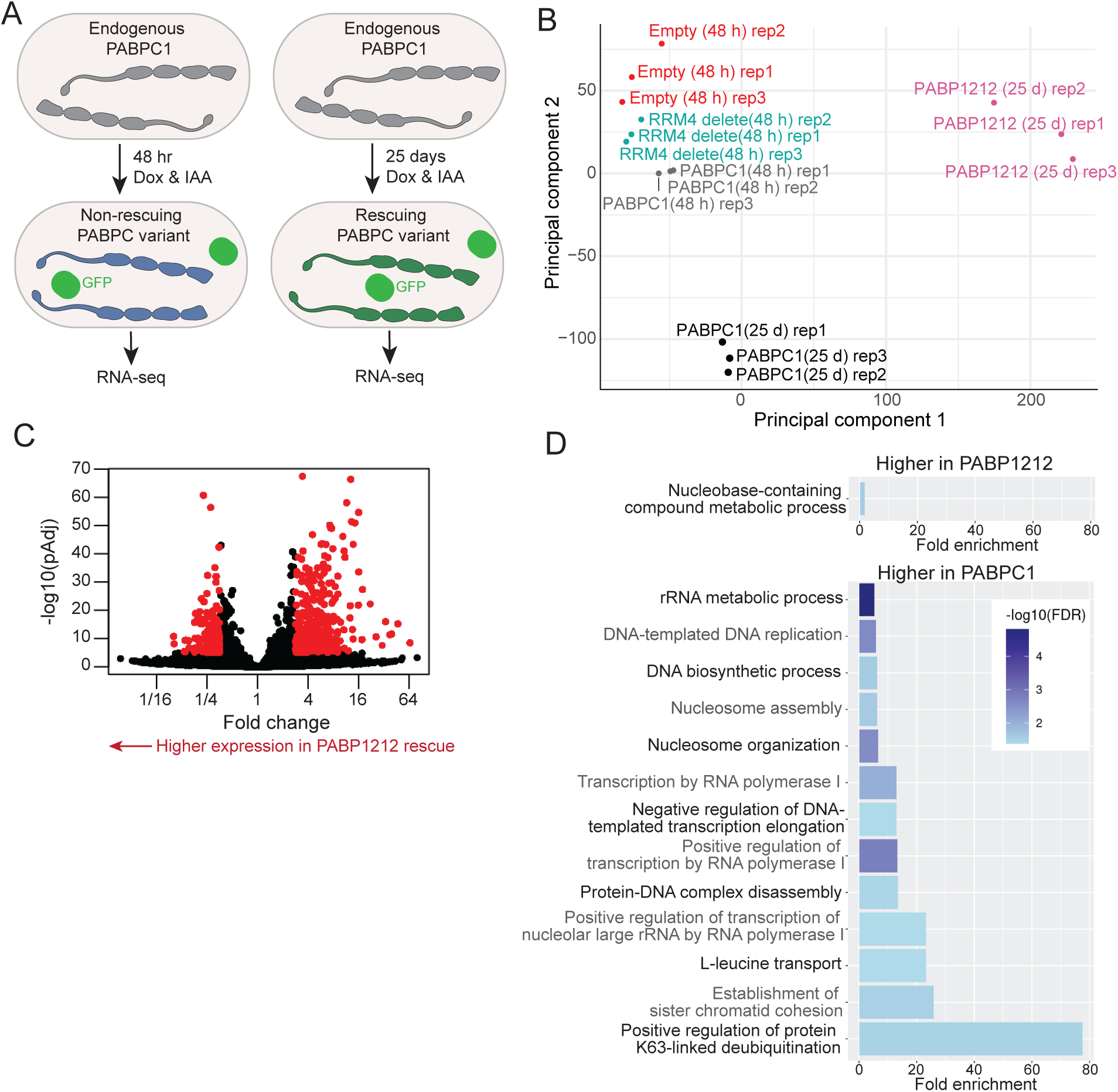
PABPC variants influence the cellular transcriptome. (A) Schematic of the experimental strategy for RNA-seq profiling. For PABPC variants that rescue, endogenous PABPC1 was replaced using doxycycline (Dox) and indole-3-acetic acid (IAA) for 25 days, followed by RNA-seq. For variants that do not rescue, endogenous PABPC1 was replaced for 48 h before RNA-seq. (B) PCA of transcriptome profiles indicating differences between PABPC1 and PABP1212 replacement. (C) Volcano plot showing the mRNA changes observed after substituting PABPC1 with PABP1212. Each dot represents a uniquely-mapped mRNA. Red points meet a cut-off of pAdj < 10^-5^ and an absolute value log2-fold change > 1.5 used for subsequent gene ontology analysis. (D) Gene Ontology (GO) term over-representation for transcripts upregulated or downregulated in PABP1212 rescue relative to wild-type PABPC1 rescue (pAdj < 10^-5^ and an absolute value log2-fold change > 1.5). Enrichments for the most enriched biological processes are plotted, with bars colored by false discovery rate (FDR).

We next compared the transcriptomes of variants that were able to rescue growth (fig S2B). Among these variants, PABPC1 clustered closest to a variant in which its RRM4 domain was replaced with the RRM4 domain of *S. cerevisiae* Pab1, whereas PABPC4 replacement, or RRM4 replacement with the RRM4 domain of PABPC1L resulted in larger transcriptome differences.

PCA revealed that transcriptome profiles from *PABP1212* rescue clustered distinctly from those of wild-type PABPC1 rescue (fig 3B). Furthermore, comparison of acute (48 h) versus long-term (25 d) depletion demonstrated pronounced time-dependent transcriptome changes, with *PABP1212* rescue maintaining a distinct expression signature over WT PABPC1 rescue for both replacement durations (fig 3B).

The distinct clustering difference between WT PABPC1 and PABP1212 prompted further investigation of differential gene expression between these two rescues, with a focus on genes with absolute value log2-fold change > 1.5 and adjusted p-values (pAdj) < 10^-5^ (fig 3C). In the *PABP1212* rescue, differentially expressed genes were significantly enriched for functions related to rRNA metabolism, DNA replication, nucleosome organization, and transcription by RNA polymerase I (Figure 3D), suggesting that specific PABPC variants might differentially influence particular sets of transcripts within these gene ontology categories.

### PABPC variant replacement can be titrated to further interrogate variant function

Given that rescue proceeded differently for WT PABPC1 and PABP1212 at both the differences in growth kinetics and the differences in the transcriptome, we wondered how rescue dosage influenced the outcome. Because rescue variants were introduced into the cell line as a polyclonal, transposon-mediated integration, with variable number of integrations to implement dosage-dependent assays, and eGFP was expressed downstream of each PABPC variants, eGFP expression served as a proxy for PABPC variant expression. Thus, eGFP fluorescence provided a quantitative metric for analysis of the effect of variant expression on growth or tail lengths (fig 4A).

**Figure 4.**
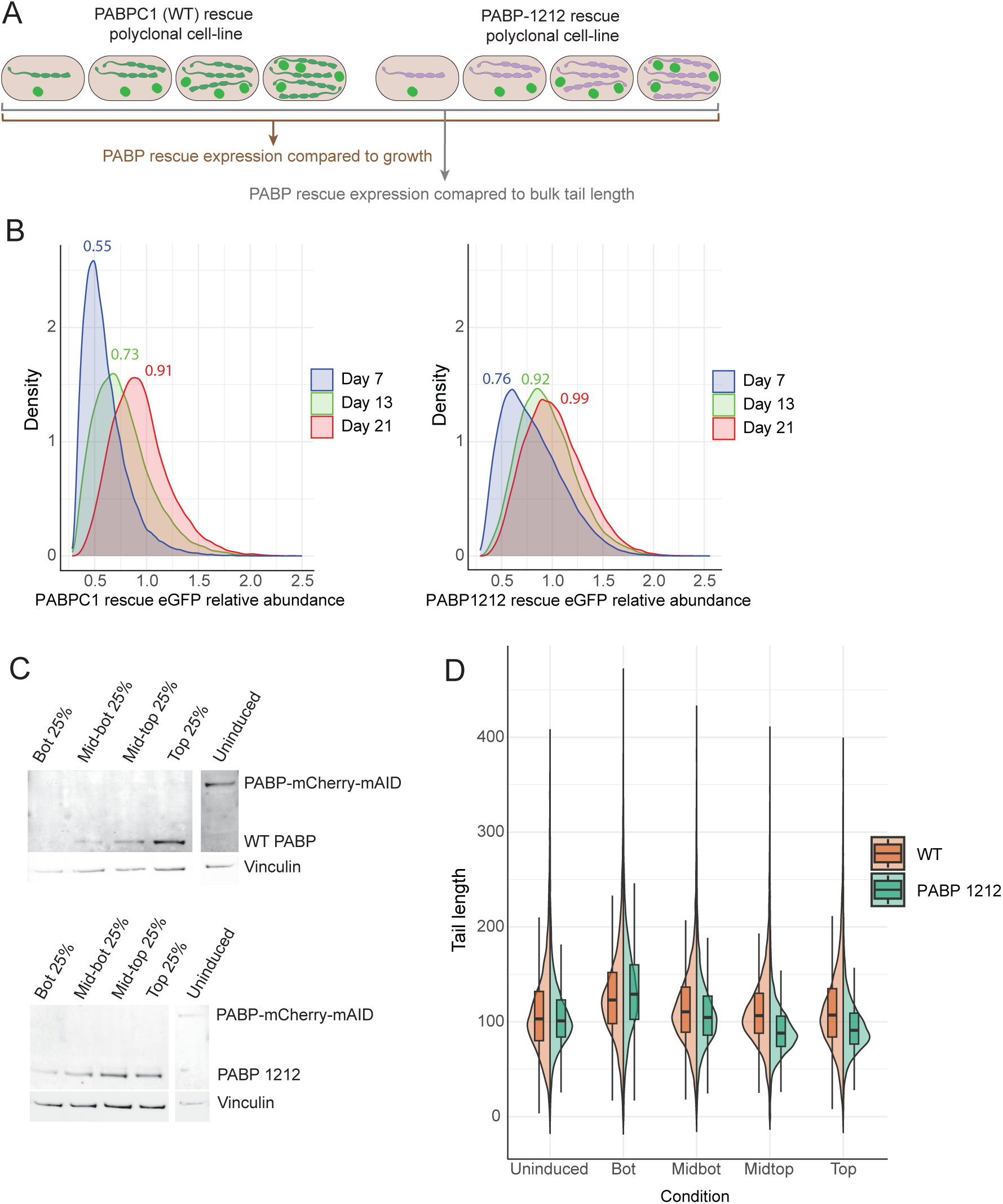
PABPC variant replacement can be titrated to further interrogate variant function. (A) Schematic of the experimental workflow for competitive growth assays, protein-abundance measurements, and tail-length measurements in polyclonal rescue lines. Endogenous PABPC1 and PABPC4 were tagged with mAID for auxin-induced degradation, and rescue variants were expressed from doxycycline-inducible constructs co-expressing GFP. (B) Competitive growth assay tracking GFP fluorescence distributions over time (Day 7, Day 13, Day 21) for (left) WT PABPC1 rescue and (right) PABP1212 rescue. Kernel density plots show shifts in GFP expression, indicating differences in competitive fitness. Median values reported for each density. (C) Immunoblot analysis of WT PABPC1, *PABP1212*, and endogenous PABPC-mCherry-mAID levels across GFP expression quartiles, with vinculin as loading control. (D) Poly(A)-tail length distributions for mRNAs of WT PABPC1 rescue and *PABP1212* rescue lines, measured in uninduced and sorted GFP-intensity quartiles (bot., midbot, midtop, and top). Tail-length distributions were obtained by nanopore sequencing and compared across GFP-based expression groups.

With respect to growth following WT PABPC1 or PABP1212 rescue, at day 7 there was lower expression for the WT PABPC1 rescue vs the PABP1212 rescue, but by day 21, both cell lines reached similar expression levels. At day 13 cells were sorted based on GFP fluorescence into roughly four quartiles, using the same fluorescence bin parameters for both WT rescue and PABP1212 rescue. Immunoblotting confirmed GFP-based quartiles corresponded to graded PABPC abundance (fig 4C). From these sorted cell populations, we acquired tail-length measurements using direct RNA nanopore sequencing (fig 4D). For both WT rescue cells, as well as PABP1212 rescue cells, lower PABPC expression resulted in longer tail lengths, consistent with the established model that limiting PABPC1 preferentially destabilizes short-tailed cytoplasmic transcripts^9^. In WT rescue cells, mean tail lengths were more consistent across GFP quartiles, whereas in PABP1212 rescue cells, mean tail lengths showed larger dynamic range across expression levels (fig 4D, S3A).

## Discussion

In this study, we establish a degron-mediated protein replacement platform that enables acute removal of endogenous PABPC1 and PABPC4 and their functional substitution with engineered PABPC variants. By pairing rapid degradation with tunable rescue expression, we create a system that can disentangle the contributions of individual domains, paralogs, and engineered modules to diverse cellular roles.

Our findings reveal several principles governing PABPC1 function in human cells. First, the essentiality of RRM4 in growth rescue, together with the inviability of mutants predicted to disrupt its RNA-binding capacity, underscores a critical requirement for robust RNA engagement. By contrast, mutations that impact post-translational modifications within RRM4 appear to have small effects in vivo, suggesting that the regulatory roles of these modifications are either subtle, context-dependent, or compensated by alternative mechanisms of PABPC regulation.

Second, our systematic RRM swap experiments highlight the domain constraints that govern PABPC function. Although several heterologous RRM4 domains, including the RRM4 of the yeast ortholog, could restore growth, others—particularly from low-abundance paralogs such as PABPC5 and PABPC4L—were unable to do so. These differences likely reflect evolutionary divergence in RNA affinity, sequence preference, or interaction surfaces, and imply that paralog-specific functions may arise from distinct contributions of RRM4. Furthermore, the ability of non-native PABPC1 RRM domains to substitute for RRM4 suggests that high-affinity RNA engagement, rather than strict sequence recognition, is the primary requirement for supporting core PABPC functions.

Third, transcriptome profiling uncovered substantial variant-specific regulatory signatures. Variants that failed to rescue growth showed widespread expression changes reminiscent of PABPC depletion, whereas rescuing variants segregated based on domain composition. The PABPC1212 construct, which imposes strong poly(A)-specific binding through duplicated RRM1 and RRM2 domains, produced a transcriptome markedly distinct from wild-type rescue. Enrichment for genes involved in rRNA metabolism, DNA replication, and chromatin organization indicates that altering PABPC’s RNA-binding footprint or sequence specificity can influence diverse regulatory networks. These findings raise the possibility that PABPC could shape gene expression not only through stabilizing short-tailed transcripts but also via broader effects on translation capacity, RNA homeostasis, or nuclear feedback pathways.

Finally, dosage-dependent rescue experiments demonstrate that PABPC abundance strongly influences poly(A) tail length in vivo. Reduced PABPC rescue expression consistently produced longer tails, aligning with models in which PABPC preferentially stabilizes short-tailed cytoplasmic mRNAs^9,15^. The heightened tail-length sensitivity observed for the PABP1212 variant suggests that distinct RNA-binding domain compositions modulate the relationship between PABPC abundance and deadenylation dynamics, with potential implications for transcriptome remodeling in contexts of stress, development, or disease.

Together, our protein replacement system provides a useful tool for dissecting the multiple functions of PABPC and, more broadly, for studying essential RNA-binding proteins that operate at high cellular concentration. Future work could extend this framework to investigate PABPC contribution to translation initiation, coupling between deadenylation and decapping, or the interplay between PABPC and translational repressors. Moreover, the tunability of this system and its compatibility with variant libraries opens the door to high-throughput domain mutagenesis, evolutionary reconstruction, and functional readout of variant protein domains. By enabling precise, rapid, and reversible replacement of essential poly(A)-binding proteins, this platform offers a foundation for broad mechanistic studies of mRNA regulation in human cells.

## Methods

### Plasmid construction

Plasmids pRM026a, pRM026c, and pRM026e-pRM026u were cloned for replacement of endogenous PABPC1 and PABPC4 with various PABPC variants. Each variant was synthesized as a gene block (IDT) or by amplifying segments of human PABPC1 with overlapping 20-nt homology arms and assembled into a dox-inducible sleeping beauty expression vector via Gibson assembly. Plasmid pRM029 was cloned into pDarmo via Gibson assembly, for expression of SBP-tagged PABPC1 under the CMV promoter. Whole plasmid sequencing using nanopore (Plasmidsaurus) confirmed each plasmid was constructed as intended.

### Cell line generation

Cell lines were generated via co-transfection of a plasmid expressing sleeping beauty transposase and transfer plasmids pRM026a, pRM026c, and pRM026e-pRM026u which each express a different PABPC variant. After a 2-day recovery, cells were then selected for transposase-mediated integration via blasticidin resistance.

### Cell culture and growth assays

HCT116 cells were cultured in McCoy’s 5A Medium supplemented with 10% FBS and incubated at 37 °C and 5% CO_2_. Cell lines were passaged by subcultivation (1:5 ratio) every 2–3 days at 70–90% confluency using 0.25% Trypsin-EDTA. To induce PABPC1 and PABPC4 degradation and expression of PABPC variants, cells were treated with 1μg/mL doxycycline and 0.5 mM Indole-3-acetic acid (IAA) unless otherwise indicated. To measure cell growth, each cell line was cultured in triplicate wells of a 12-well dish. At each time point, cells were washed with PBS, trypsonized from the plate, quenched with media, and counted by hemocytometer on a Countess cell counter according to manufacturer instructions.

### Determination of PABPC1 post-translational modification

Plasmid pRM029 was transfected into one well of a 6-well plate of confluent HEK cells. Cells were washed with PBS, lysed on ice, and pelleted at 20,000 x g for 15 minutes to remove cell debris. SBP-tagged PABPC1 was captured from the supernatant using streptavidin beads (Pierce) and eluted with biotin. The elution was run on a Coomassie gel and the size corresponding to PABPC1 was cut out and submitted to the Whitehead Quantitative Proteomics core for identification of post-translational modifications. The core performed both a tryptic digest and a chymotrypsin/GluC digest and ran each in DDA mode on an Orbitrap Exploris 480.

### Transcriptomic clustering

Following PABPC replacement, RNA-seq was performed on each cell line in triplicate. Total RNA was phenol chloroform extracted and 1μg RNA from each sample was used as an input for the KAPA RNA HyperPrep Kit with RiboErase. Prepared libraries were submitted for 50×50 paired-end sequencing on the NovaSeq S1, aiming for approximately 20 million reads per sample. Reads were mapped to hg38 using Spliced Transcripts Alignment to a Reference (STAR)^16^. Principle component analysis was implemented using the built-in stats package in R.

### PABPC titration by FACS

Following PABPC replacement, titration of PABPC abundance was achieved through fluorescence-assisted cell sorting (FACS). Four even gates were drawn, aiming for 25% of the population falling into each GFP expression level. A subset of the sorted cells from each population were lysed and PABPC abundance measured by immunoblotting. RNA was extracted from the remaining sorted cells by phenol chloroform extraction and total RNA was used for downstream poly(A) tail measurements by nanopore sequencing.

### Tail length measurements of PABPC replaced cells

Nanopore sequencing was used to measure transcriptome-wide tail lengths in PABP-replaced cells. 1μg total RNA from PABPC-titrated cells was used as the input for direct RNA sequencing. Sample libraries were prepared according to manufacturer instructions (Oxford Nanopore direct RNA kit SQK-RNA004) with one specific adaptation. The first ligation set was modified to instead ligate a set of custom adapters that would allow for sample barcoding and demultiplexing. Thirteen custom adapters were designed as previously described ^17,18^ and ligated onto transcript ends from each sample. Samples were sequencing using the Nanopore PromethION. Samples were demultiplexed with a custom trained model.

A Support Vector Classifier (SVC) model was constructed following previously described methods^17, 18^ to classify thirteen barcodes. Briefly, thirteen different DNA templates were transcribed *in vitro* using T7 polymerase, and each of them was ligated to a barcoded adapter, generating unique IVT transcript-barcode pairs. IVT reads from sequencing run PAY96494 were used for training, with approximately 4000 reads per label.

Ground-truth labels were established by mapping IVT transcripts to reference sequences using Minimap2^19^ for long IVT sequences and Bowtie2^20^ for short IVT sequences. Barcode identity was then inferred from the transcript sequence to which each barcode was uniquely paired during library construction. The electric current magnitude and dwell time of the barcode region in each read, termed “fingerprint”, was captured using WarpDemuX^18^. A Dynamic Time Warping (DTW)-based kernel was learned from the barcode fingerprints, and used, together with the ground-truth labels, to train the SVC.

The SVC outputs the probability that a given read belongs to each of the thirteen classes. The predicted label was assigned as the class with the highest probability, and the confidence score was defined as the difference between the highest and second-highest probabilities. The model was tested on an independently sequenced IVT dataset (run PBC20019). Confidence score thresholds for each barcode were determined using the test set to balance precision and recall. The trained model was then applied to RNA-seq data for demultiplexing, and tail lengths of the demultiplexed reads were calculated using Dorado (v0.9.5, Oxford Nanopore Technologies).

Sequencing data generated during this study will be available at GEO [Accession Number Pending].

---

**Figure S1.**
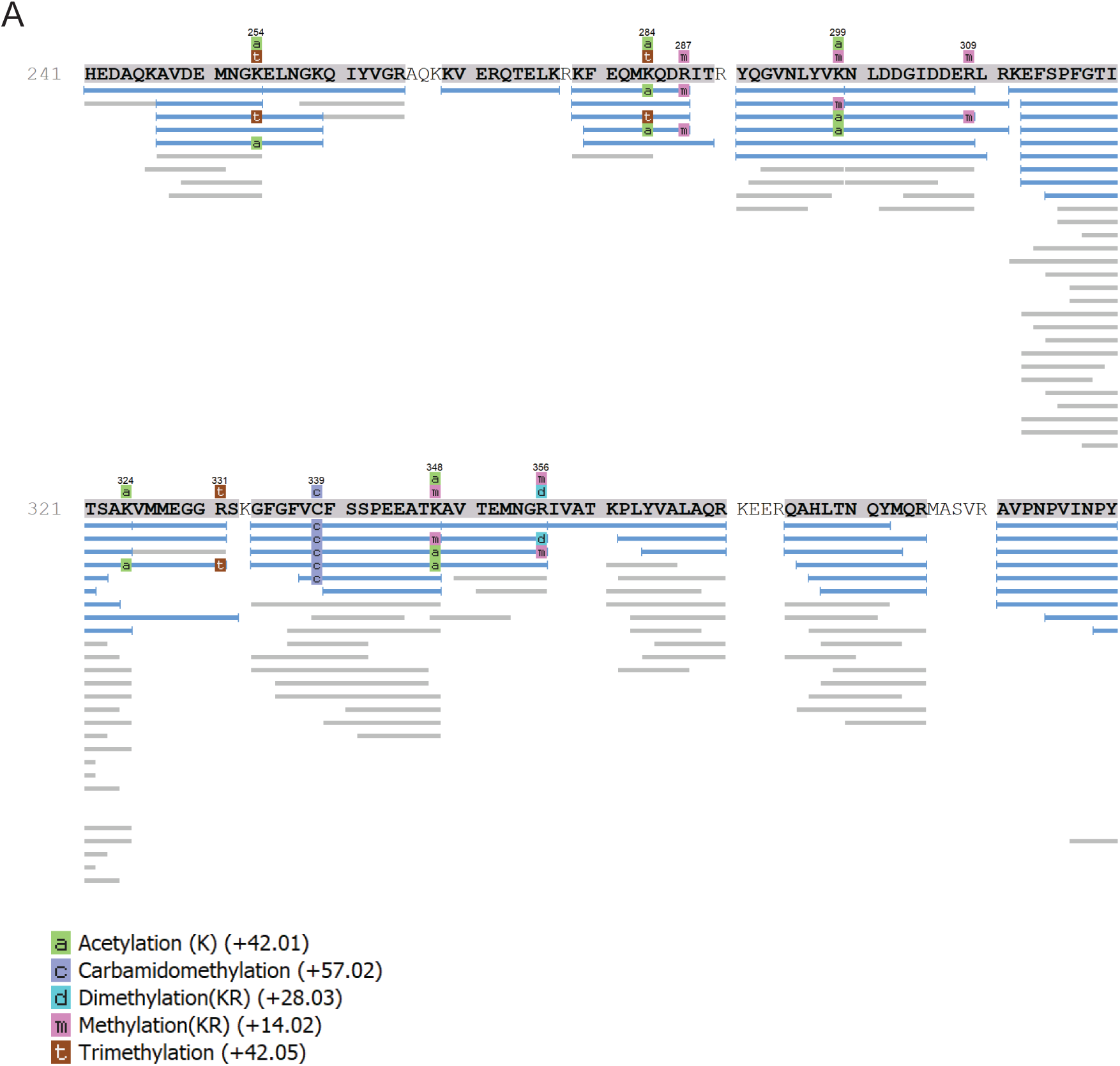
Assessment of native post-translational modifications within RRM4 of PABPC1. (A) PTM profile showing amino acid residues with observed acetylation, carbmidomethylation, demethylation, methylation, and trimethylation.

**Figure S2.**
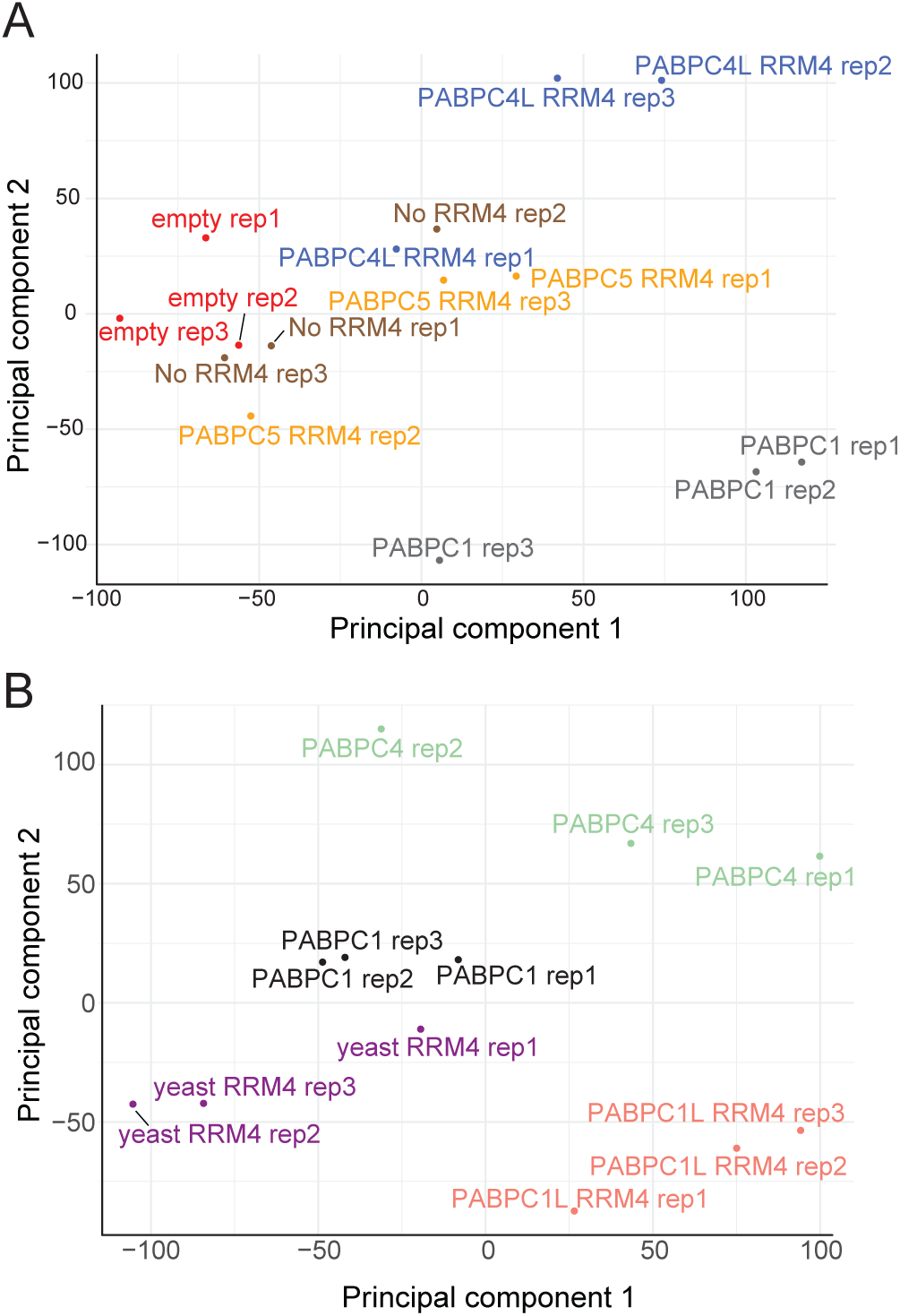
PCA comparing transcriptomes following PABPC variant replacement. **(A)** PCA of transcriptome profiles 48 h post drug treatment. **(B)** PCA of transcriptome profiles 25 days post drug treatment.

**Figure S3.**
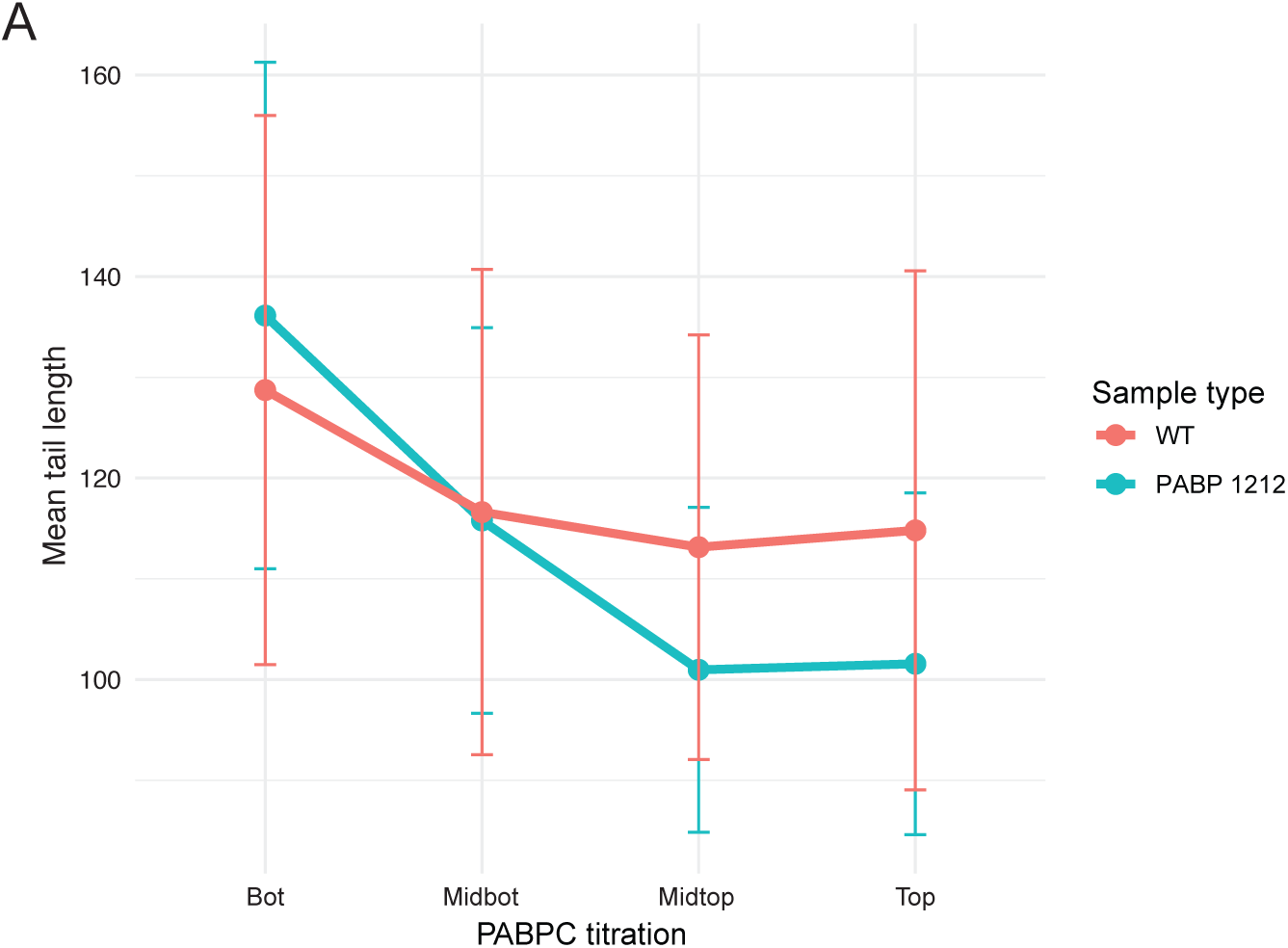
Distribution of poly(A) tail lengths following PABPC variant replacement. **(A)** Mean tail lengths plotted across PABPC titration with error bars indicating standard deviations.

## Acknowledgements

We thank the Whitehead Institute Genome Technology Core for high-throughput sequencing. We thank the Whitehead Institute Proteomics Core for PTM-aware protein mass spectrometry. R.Y.M. was a Howard Hughes Medical Institute fellow of the Damon Runyon Cancer Research Foundation (grant no. DRG-2485-23). D.P.B. is an investigator of the Howard Hughes Medical Institute. We thank A. Balabaki, B. Wierbowski, K. Xiang, A. Zhou, and other members of the Bartel lab for fruitful discussions.

## Notes

### Competing Interest Statement

The authors have declared no competing interest.

